# Distractor-response binding influences visual search

**DOI:** 10.1101/2024.03.13.584838

**Authors:** Fredrik Allenmark, Hao Yu, Hermann J. Müller, Zhuanghua Shi, Christian Frings

## Abstract

Intertrial priming effects in visual search and action control suggest the involvement of binding and retrieval processes. However, the role of distractor-response binding (DRB) in visual search has been largely overlooked, and the specific processing stage within the functional architecture of attentional guidance where the DRB occurs remains unclear. To address these gaps, we implemented two search tasks, where participants responded based on a separate feature from the one defining the target. We kept the target dimension consistent across trials while varying the color and shape of the distractor. Moreover, we either repeated or randomized the target position in different sessions. Our results revealed a pronounced response priming, a difference between trials where the response changed vs. repeated: they were stronger when distractor features or the target position were repeated than they varied. Furthermore, the distractor feature priming, a difference between the distractor features repetition and switch, was contingent on the target position, suggesting that DRB likely operates at late stages of target identification and response selection. These insights affirm the presence of DRB during visual search and support the framework of binding and retrieval in action control as a basis for observed intertrial priming effects related to distractor features.

**Public significance statement:** This study investigated inter-trial effects within visual search tasks and uncovered significant evidence for the role of distractor-response binding. This phenomenon involves linking a response in a given trial to the perceptual features (e.g. color and shape) of non-target items. Crucially, the study revealed that this distractor-response binding effect depends on whether the target location is repeated nearly repeated, suggesting that the retrieval of a previous response occurs at the later stages of target identification or response selection, even though non-target features likely are identified at an earlier stage.

## Introduction

Studying human behavior in a laboratory setting has led to fragmentation of cognitive research into paradigm-specific approaches, using strict-controlled experimental paradigms, that may not accurately reflect real-world behavior as a whole but only a certain aspect of behavior. While controlling surroundings and isolating certain aspects of behavior can help identify and pinpoint underlying processes, it can also be risky as other ‘unintended’ (from the viewpoint of the researcher using a particular paradigm) processes may be overlooked, leading to misinterpretation of results. What is more problematic is that at theoretical levels the interpretations and conclusions may fall short in fully capturing the complexities of human behavior as only parts of behavior are looked upon at a time. In this article, we present such an instance of overlooking as we argue that in established intertrial priming in compound search tasks (V. Maljkovic and Nakayama 1994; Vera Maljkovic and Nakayama 1996; Olivers and Meeter 2006), i.e. search tasks where the target is defined based on one visual feature (e.g. the target has a unique color or shape) and the response is made based on a different feature (e.g. the orientation of a line inside the target shape), other processes in the form of binding and retrieval (Frings et al. 2020) are also at work and contribute to the observed results (Henson et al. 2014). Our study is thus not just an empirical contribution, but also a call for a more integrated and holistic approach to visual search and action control theorizing (Frings et al. 2020; Lamy, Yashar, and Ruderman 2010; Lamy, D., Frings, C., Liesefeld, HR 2023; Heinrich René Liesefeld et al. 2019; Heinrich R. Liesefeld and Müller 2020; Yashar and Lamy 2011; Yashar, Makovski, and Lamy 2013). These two research strands developed mostly independently of each other (Lamy, D., Frings, C., Liesefeld, HR 2023) while we argue that they might actually profit from relating to each other and further that it is important to connect these research strands for the ultimate goal, namely to understand human behavior not only in a particular experimental paradigm but in general. Our contribution in this article could be seen as an important step in this direction.

The paradigms we connect here are still artificial in that they are established experimental paradigms run in the laboratory - so one may ask how our approach actually contributes to a more holistic understanding of human behavior in general (even outside the laboratory). For instance, action control paradigms often use simple responses as key presses while the results are interpreted in terms of actions (including quite different types of actions like playing tennis). Yet, the cognitive part of an action, the planning and matching of goals with perceptions, is actually quite comparable between a key press and a more naturalistic movement (say hitting a tennis ball with your racquet) while the action execution part is obviously quite different. Our approach here is concerned with paradigms as used in the event-coding and action control literature (Hommel 2004; Frings et al. 2020) that traditionally focus on the planning part of an action. The same holds true for visual search tasks - the search aspect in the laboratory might be actually quite comparable to searching in the real world (for a review, see Wolfe 2021). Against the background of these considerations, bridging visual search and action control studies in the laboratory might be a promising road to approach understanding human behavior more generally.

Intertrial priming is a phenomenon where the outcome of a previous trial influences the performance and processing of subsequent trials in a visual search task. In a seminal series of studies, Maljkovic & Nakayama (1994; 1996, 2000) have shown that search is much faster when the target’s color or location repeats on successive trials (*intertrial priming of feature and location*, respectively). In addition, repeating the color of distractor items can also facilitate search processes (see also Lamy et al. 2008), a phenomenon termed as *distractor-repetition priming*. Similarly, Müller and colleagues (Found and Müller 1996; Müller, Heller, and Ziegler 1995; Müller, Reimann, and Krummenacher 2003) observed a structurally similar effect. Participants searched for a singleton target that could pop out from the surrounding nontargets by either its unique color or its unique orientation; the specific target feature (e.g., red vs. blue) as well as the feature dimension (color vs. orientation) unpredictably changed across trials. Results indicated a strong performance advantage when the target dimension repeated vs. switched. Müller and colleagues proposed the Dimension Weighting Account (DWA) of visual attention (for reviews, see Krummenacher and Müller 2012; Heinrich René Liesefeld et al. 2019; see also Heinrich R. Liesefeld and Müller 2020) to account for inter-trial dimension priming. The DWA assumes that signals from the target dimensions are up-weighted, while those from the distractor dimensions are down-weighted. Selecting a pop-out target on a given dimension on Trial *n* increases the weight of that dimension, an increase that typically persists until (at least) the next trial (Allenmark, Müller, and Shi 2018). This weight adjustment enhances target guidance in the next trial if it shares the same dimension, partially accounting for dimension-specific intertrial effects (Müller and Krummenacher 2006)^1^.

The DWA framework has evolved into the multiple weighting-systems (MWS) account (Zehetleitner, Rangelov, and Müller 2012; Rangelov, Müller, and Zehetleitner 2012), which assumes that “processing at multiple stages (e.g., target selection, response selection) on any given trial *n* can lead to the laying down of separate, stage-specific memory traces that in turn influence the processing on the respective stages on the subsequent trial *n+1*” (Zehetleitner, Rangelov, and Müller 2012, 879). This concept is particularly evident in compound search tasks (e.g. Zehetleitner, Rangelov, and Müller 2012), where participants first search for a target based on a specific feature (e.g., color) and then make a discrimination response based on another feature (e.g., orientation). In those tasks, the intertrial priming is structurally equivalent to sequential priming tasks used in the action control literature (Frings et al. 2020). Zehetleiter and colleagues, for instance, observed an interaction of response and color dimension. Specifically, participants performed better when both the response-defining and target-defining features were repeated compared to trials where only one of these aspects was repeated. In the *d*istractor-*r*esponse *b*inding paradigm (DRB; Frings, Rothermund, and Wentura 2007), an analogue interaction is observed. In a typical distractor-response binding task, participants focus on the target while ignoring the flank distractors in a letter array. Performance improves when both the response and an irrelevant distractor from the previous trial (*n-1*) are repeated in the current trial (*n*). However, the interpretation of DRB differs significantly. The DRB paradigm, a well-established task in the field of action control, assumes the integration or binding of all features, including the response, from the previous trial into an event-file (Hommel 2004, 2005; Frings et al. 2020). When any feature from the previous trial is repeated, the entire previous event-file including response features, is retrieved, influencing the response of the current trial. This Binding and Retrieval in Action Control (BRAC) approach has been extensively validated across various settings, stimuli, modalities, and laboratories (for a review, see Frings et al. 2020). DRB remains central to current theories in action control, with a general consensus that binding and retrieval processes broadly contribute to action (Hommel 2004; Frings et al. 2020; Kiesel et al. 2023).

The concept of retrieval has already been employed to explain certain inter-trial priming effects in earlier studies (Huang, Holcombe, and Pashler 2004; Ásgeirsson and Kristjánsson 2011; Yashar and Lamy 2011). For instance, Huang and colleagues (2004) have demonstrated that repeating task-irrelevant target features, such as color in an orientation judgment task, can facilitate search performance. They suggested that “these irrelevant features belonged to the target objects. Attention to an item - in this case, the target of the search - may automatically trigger processing of all its features, including irrelevant ones” through episodic memory traces (Huang, Holcombe, and Pashler 2004, 18). Following this, many other studies investigated how task-irrelevant target features affect search performance, and finding that inter-trial priming based on episodic memory typically occurs in difficult searches (Ásgeirsson and Kristjánsson 2011; Lamy, Zivony, and Yashar 2011) and mainly at the late response selection stage (Lamy, Yashar, and Ruderman 2010; Zehetleitner, Rangelov, and Müller 2012). However, what has been largely neglected is the role of the distractor-response binding in visual search (but see Lamy et al. 2008). Research on the influence of episodic retrieval has predominantly focused on the manipulation of irrelevant features of the target, based on the implicit assumption that selecting a target also activates its related features. This ‘target-centered’ episodic memory perspective differs fundamentally from the response binding account. The latter proposes that all features, including those from distractors, can be linked to the response, thereby affecting subsequent responses.

### The present study

Considering the structural resemblance between intertrial effects in visual search and action control, we propose that effects of binding and retrieval also operate in compound visual search tasks, although past studies may have overlooked stimulus-response bindings. Our study has two main objectives: first, to explore whether the distractor-response binding (DRB) effect seen in the flanker task can also be manifested in standard visual search tasks; and second, to pinpoint the stage within the attentional guidance architecture where DRB occurs. A distinct difference between the flanker task and the search task is the inclusion of a search stage in the latter. We hypothesize that if DRB occurs during the early search stage, it would be more effective in suppressing irrelevant distractors and facilitating target localization. Given that the distractors usually outnumber the target in a display, suppressing repeated distractors would take place at a preattentive stage, before the target is located. Hence, DRB facilitation should be independent of the target position. In contrast, if DRB occurs at the later stages of target identification and response selection, we expect DRB to be influenced by the target’s location and to be activated particularly when the target is near its previous position.

To test these hypotheses, we adjusted the compound task to keep the target dimension constant, avoiding potential confounding issues related to target-related dimension-weighting facilitation. Specifically, each trial had a consistent target, but with varied distractor color and response. The target was redundantly defined by both color and shape, remaining constant while the distractors always differed in color and shape from the target and varied randomly across trials. Participants were instructed to respond based on a different feature of the target (orientation in Experiment 1 and a black dot’s location in Experiment 2). This approach fosters a feature-based target template rather than a dimension-specific guided search. Consequently, according to DWA, the effect, if any, would be minor^2^. In contrast, the BRAC model predicts impacts of distractor and response repetition. Specifically, it would predict that the effect of response repetition (the response priming effect) should be larger when the same distractor features are repeated, compared to when the distractor features change from one trial to the next. For partial repetitions, the prediction of the BRAC model would depend on what strategy participants use to locate the target. If participants rely equally on both features, BRAC would predict that the response priming effect on partial repetition trials should be intermediate between the effect on full repetition and full change trials. However, if participants learn to rely mainly on one feature for target identification, because this feature is more salient or more task-relevant, BRAC would predict that the response priming should be mainly influenced by that feature with less or no dependence on the other feature. To mimic the non-search flanker task, we fixed the target position in one session, while letting the target position change at random in another session. If DRB takes place at an early search stage, we expect distractor feature priming in both cases. In contrast, if it occurs later, during target identification or response selection, distractor feature priming should only appear when the target position is repeated or near its previous location.

To foreshadow the results, in two experiments (N = 24) using typical search displays, we found both the repetition of distractor features and the target position influenced response priming. Response priming was most noticeable when the target was repeated or near the previous target location, while distractor feature priming was mainly shown in the response repeated trials. The results reveal a strong response binding effect in typical visual search, which occurs at the late stage of response selection.

## Experiment 1

In Experiment 1, participants searched for a green ellipse target and indicated its orientation - either clockwise or counter-clockwise. The target’s shape and color remained consistent throughout all trials, while the distractors varied shape and color, independent of the response feature of the target. According to the DRB, we would expect an interaction between the distractor feature and the response, which would closely resemble an intertrial priming effect.

### Methods

#### Participants

24 healthy individuals, normal or corrected-to-normal visual acuity, and normal color perception passed with the Ishihara color test (Clark 1924), were recruited for Experiment 1 (mean age ± SD: 26.9 ± 5.54 years; age range: 22–43 years; 15 females). The sample size was determined based on past DRB studies that usually found medium to large effect sizes (dz between 0.4 and 1; e.g., Frings and Moeller 2010, 2012; Koch, Frings, and Schuch 2018; Moeller and Frings 2011). To achieve a power of 80% (1-ß) with a median effect (*d_z_* = 0.6), at least 19 participants were needed. To balance all factors, we determined 24 participants for the present study. All participants provided written informed consent prior to the experiment and were compensated their participants with 9 Euros per hour or course credit. This study was approved by the ethics board at the LMU Faculty of Pedagogics & Psychology. All data in Experiment 1 was collected in 2021.

#### Apparatus and Stimuli

The experiment was conducted in a sound-attenuated and moderately lit test room. Participants sat in front of an LCD display monitor with a resolution of 1920 × 1080 and a refresh rate of 120 Hz, with a viewing distance of 57 cm with the aid of a chinrest. The experiment was created in PsychoPy (version 2022.1.3). The visual search items were 22 shapes arranged on three concentric circles, with four on the innermost circle (eccentricity of 3°), eight on the middle circle (eccentricity of 6°), and ten on the outer circle (eccentricity of 9°). The search target, which had a unique shape and color - a green (CIE [Yxy]: [33.5, 0.240, 0.397]) ellipse (2.3° × 1.1°), could appear at any position on the middle circle. The non-targets, however, varied in both color and shape from trial to trial (but were homogeneous on any given trial). The possible non-targets could be brown (CIE [Yxy]: [26.1, 0.378, 0.376]) or blue (CIE [Yxy]: [31.8, 0.2000, 0.265]) rectangles (2.3° × 0.6°) or diamonds (2.3° × 1.8°) (see Figure 1). All shapes were randomly tilted 3 degrees either clockwise or counter-clockwise from the vertical orientation, and the search task was to discriminate the orientation of the target.

**Figure 1.**
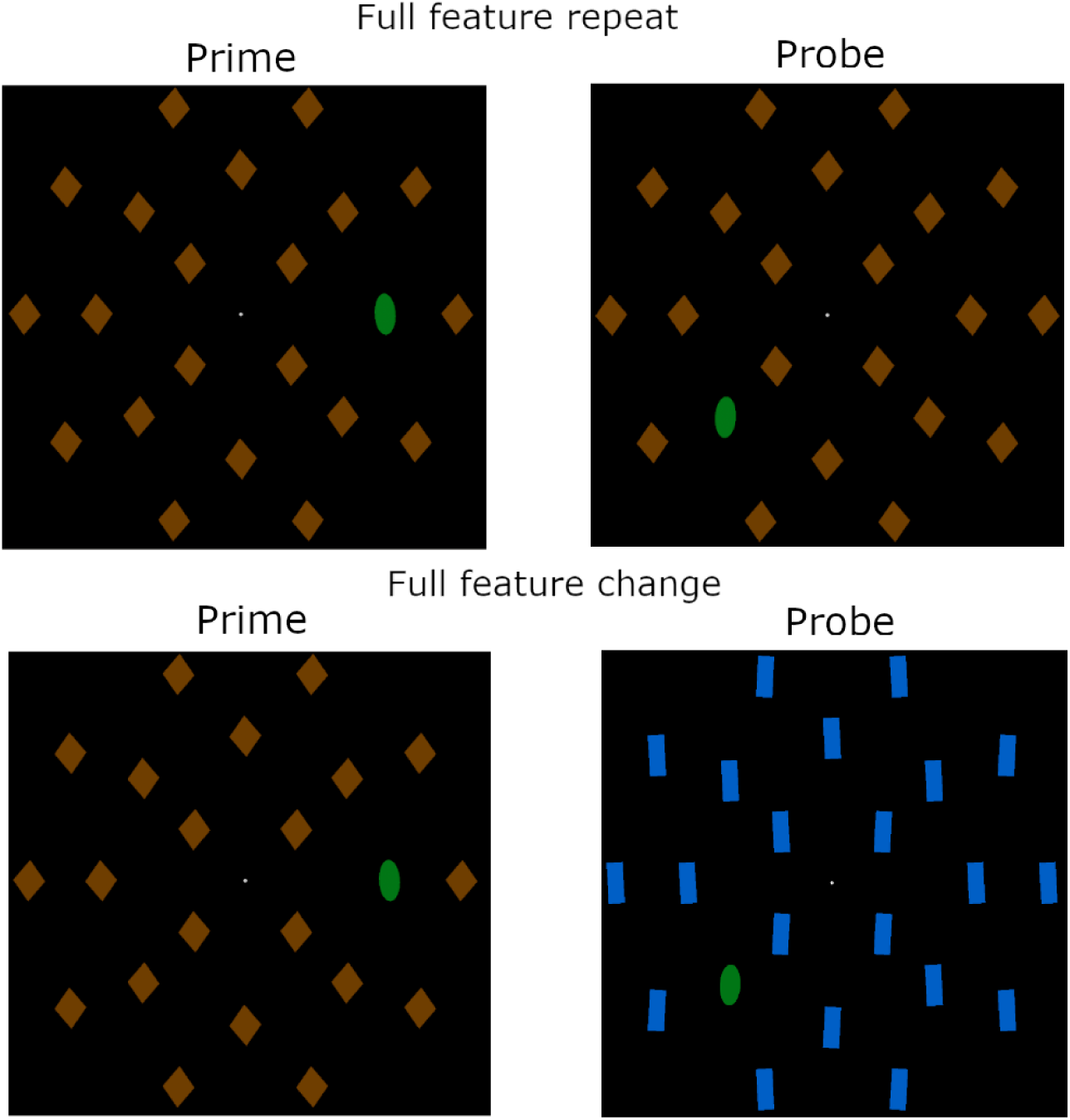
Examples of search displays used in Experiment 1. In the full feature repeat condition, the non-target color and shape were repeated between the prime and probe trial, while they both changed in the full feature change condition. The task was to find the green ellipse target, which was unique in both shape and color, and discriminate its orientation clockwise or counter-clockwise.

#### Design and procedure

The experiment consisted of 16 mini-blocks of 128 trials each. Each mini-block had pairs of trials, where the first was called “prime” trial and the second the “probe” trial. This method is commonly used in the action control studies to have a better control of binding and retrieval (Moeller and Frings 2022). Participants were required to press the ‘space’ button to start the next pair of trials. The non-targets were either brown or blue and in the shape of a diamond or rectangle, while the target was a tilted left or right ellipse. The distractor features and the target orientation could either repeat or change from the prime to probe trial. Each of the eight possible combinations occurred an equal number of times in the prime-probe trial pairs. The 16 mini-blocks were divided into two sessions, “fixed location” and “random location”, both comprising eight mini-blocks. In the “fixed location” session, the target remained in the same position within each mini-block but changed randomly between mini-blocks. In the “random location” session, the target’s location changed randomly from trial to trial, including between the prime and the probe trial in each pair. The order of the two sessions was counterbalanced among participants.

Each trial began with a fixation cross for a random duration of 700 and 1000 ms. Following the fixation period, the search display appeared and remained on the screen until a response was made. The task was to find the target and indicate its orientation as soon as possible by pressing either ‘s’ or ‘l’ key, representing counter-clockwise and clockwise, respectively. An error feedback display with the message “Incorrect” was presented for 500 ms after an incorrect response, while no feedback was given for correct responses. After a prime trial, the fixation period of the next (probe) trial followed directly, while probe trials were followed by a self-paced break, which could be ended by pressing the ‘space’ button.

#### Transparency and openness

The experimental code, raw data, and data analyses of the present study are publicly available at: https://github.com/msenselab/distractor_binding

This study was not preregistered.

### Results

In this study, we focused on two primary effects: distractor feature priming and response priming. We assessed these effects by evaluating the priming in prime-probe pairs, specifically between conditions of full repetition versus full change, and by comparing the performance in probe trials. Specifically, distractor feature priming refers to the performance difference observed between trials with fully changed distractor features and those with identical repeated distractor features. We used this measure to examine the effect of distractor-binding. Similarly, the response priming effect measures the performance difference in trials where the response changed versus repeated, which is a key measure in exploring the response-binding effect. For thoroughness, we also included the analysis of partial distractor repetition in the Appendix.

We concentrated on those pairs where both responses were correct. On average, there were about 8.2% of error trials, where the error rates (ERs) on the probe trials varied between 2% to 6% (see Figure A1 in the Appendix). In addition, we removed those outliers where response times (RTs) were slower than 3 s or faster than 200 ms (less than 1%). We observed a similar trend in RTs and ERs (see the Appendix) - slower RTs with higher error rates, which ruled out any potential speed-accuracy trade-off. To better compare across different conditions, we used the inverse efficiency score (Townsend and Ashby 1983), IES = RT/(1-ER), to measure performance (see the Appendix for the separate reaction time and error rate results, and for an analysis of partial distractor feature repetition trials).

Figure 2 shows the response priming effect (A) and the feature priming effect (B) for different conditions. A repeated-measures ANOVA on the response priming effect with the factors of Distractor Feature (full repetition vs. full change) and Target Position (fixed vs. random) revealed that both main factors were significant: Distractor Feature, *F*(1, 23) = 7.25, *p* = .013, *η*_*p*_^2^ = .24, this is the test the power-analysis was calculated for., and Target Position, Distractor Feature, *F*(1, 23) = 7.25, *p* = .013, *η*_*p*_^2^ = .24. However, the interaction between Distractor Feature and Target Position was not significant, *F*(1, 23) = 0.27, *p* = .61, *η*_*p*_^2^ = .01.

**Figure 2:**
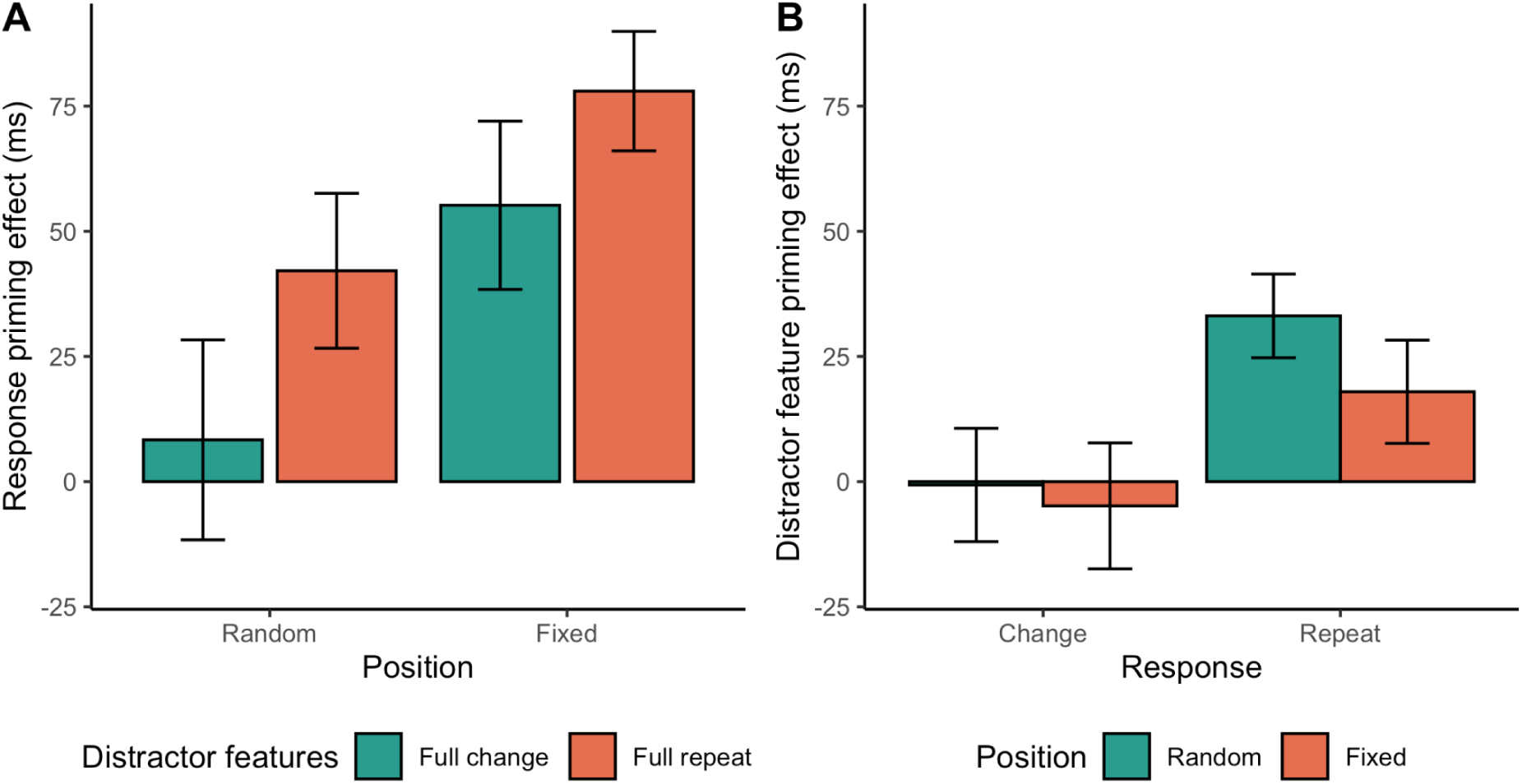
Response priming effect (A) and distractor feature priming effects (B) measured by the inverse efficiency score (ms) from the probe trials, separated by target location block (Random vs. Fixed) and the distractor feature (Full repetition vs. Full change) or response repetition in Experiment 1. Error bars indicate the standard error of the mean.

The response priming effect showed a 28 ms increase for trials with full distractor repetition relative to those with full change. Further analysis, including partial repetition (reported in the Appendix), showed that the response priming was mainly driven by the change vs. repetition of the distractor shape, rather than the distractor color. This suggests that the shape and color dimensions did not equally contribute to the response priming, partly owing to their relevance to the task we used here - the orientation discrimination of the target. The response priming was more marked (41 ms larger) for the fixed relative to the random target location. Critically, the response priming for the full-changed distractor features with random target positions was not different from zero, *t*(23) = 0.43, *p* = .67, indicating changing all features would diminish response priming. Thus, we observed both the distractor feature and the target location influence response priming.

Conversely, the distractor feature priming shows a different pattern (Figure 2B). The effect of Distractor Feature was generally smaller than the response priming (11 ms vs 46 ms). It appeared primarily in trials where the response was repeated (26 ms on response repeated trials vs. -3 ms on response change trials). A repeated measures ANOVA on the distractor feature priming, considering Response Repetition (repeat vs. change) and Target Position (fixed vs. random), revealed a significant effect of Response Repetition, *F*(1,23) = 6.0, *p* = .022, *η*_*p*_^2^ = 0.21. However, there was no significant effect for Target Position, *F*(1,23) = 0.92, *p* = .35, *η*_*p*_^2^ = 0.04), nor for their interaction, *F*(1,23) = 0.27, *p* = .61, *η*_*p*_^2^ = 0.01. The distractor feature priming effect was significantly greater than zero in response-repeated trials (*t*(23) = 3.3, *p* = .0029), but not significantly different from zero in response-changed trials (*t*(23) = -0.37, *p* = .72). Since we observed a significant distractor feature priming effect only when the response was repeated, it could be a result of retrieval of the correct response when both the distractor features and the response were repeated.

Given the target position played a critical role in the response priming effect, we further investigated the target distance between the prime and the probe pair in the random target session, which showed how the prime-probe target distance influences the response priming (see Figure 3). There was a significant effect of the target distance, *F*(2.6,59.6) = 4.23, *p* = .012, *η*_*p*_^2^ = .16, showing that the response priming effect tended to be smaller as the target moved a greater distance. We performed an additional post hoc analysis, in order to examine for which target distances the response priming effect was significantly greater than zero, which revealed that the response priming effect was only significant when the target was repeated at the same position and marginally significant when the target moved to a neighboring location (target position repeated: *t*(23) = 3.05, *p*_bonf_ = .028, target position at the neighbor of the previous target position: *t*(23) = 2.77, *p*_bonf_ = .055, all other *p*s > 0.15). However, there was no significant interaction with the repetition of the distractor features, *F*(2.4,55.8) = 1.47, *p* = .24, *η_*p*_*^2^ = .06, but there was a significant main effect of feature repetition (*F*(1,23) = 4.96, p = .036, *η*_*p*_^2^ = .18).

**Figure 3:**
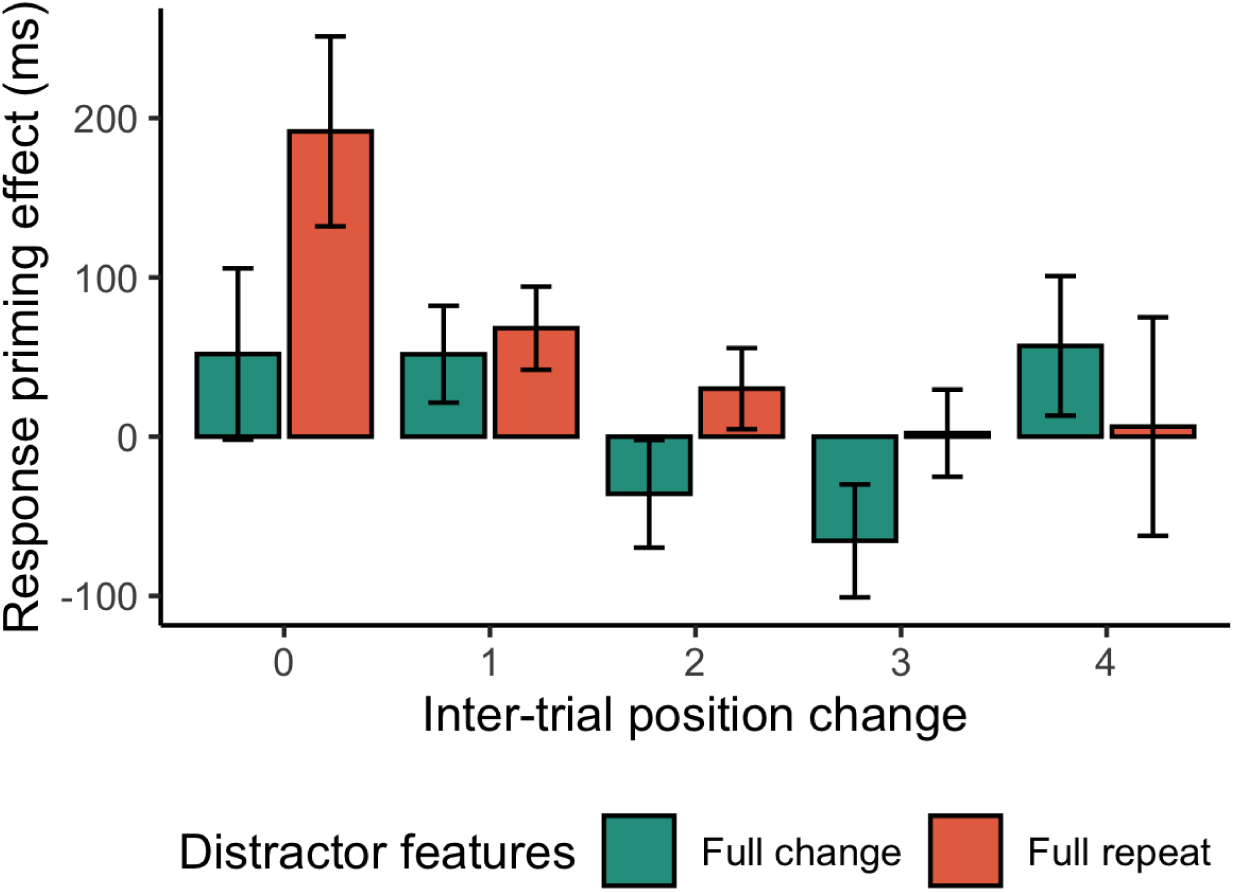
Response priming effect on the inverted efficiency score (IES), in the random target location block of Experiment 1, as a function of the distance the target moved between prime and probe trial (inter-trial position change). Error bars indicate the standard error of the mean. The significant response priming effect was observed when the target position and the distractor features were repeated (the single-out red bar, *t*(23) = 3.05, *p*_bonf_ = .028).

### Discussion

By keeping the target features constant but varying the distractor features, we revealed both the response priming effect and the distractor feature priming effect. The response priming was more marked when the distractor features were fully repeated compared to the fully changed, and when the target position was fixed relative to change. These findings are in line with the prediction of the DRB account (Frings, Rothermund, and Wentura 2007). The distractor feature priming was similar to the previous report (Lamy et al. 2008). When the distractor features were fully repeated, relative to fully changed, the performance was facilitated.

Further analysis of the response priming in the random target position session revealed that the response priming was mainly contributed by the condition when the target position was repeated. This suggests that the response priming is unlikely to have originated during the early search stage where the target location has not yet been identified.

While the results are in good agreement with the distractor-response binding account (Frings, Rothermund, and Wentura 2007), one might argue that the response priming could be attributed, at least partially, to the repetition of the target features. Specifically, the target features (i.e., a tilted ellipse) were coupled with the response, implying the response repetition was also a repetition of the target feature. The distinct combination of target features (elliptic shape and orientation) might pop out, and the orientation can be processed without focused attention, which subsequently promotes the adoption of the target feature searching strategy. In order to address this potential confound, in Experiment 2 we used a more spatially localized feature, a small black dot inside a circular target, as the response-defining feature, rather than the orientation of the entire target. By making the response-defining feature more spatially localized, and thereby increasing the need to move the target into focal attention in order to make the necessary discrimination, we aimed to limit any preattentive feature processing that may promote the target feature search strategy.

## Experiment 2

### Methods

#### Participants

24 healthy participants, with normal or corrected-to-normal visual acuity, and normal color perception, were recruited for Experiment 2 (mean age ± SD: 22.5 ± 2.9 years; age range: 19–29 years; 20 females). All provided written informed consent prior to the experiment and were compensated their participants with 10 Euros per hour or course credit. This study followed the ethical guidelines of the local ethic committee at Trier University and was in accordance with the recommendations of the German Psychology Association. All data in Experiment 2 was collected in 2022.

#### Apparatus and Stimuli

The setups were similar to those used in Experiment 1, but with two important differences. First, the target and non-target shapes both had equal width and height. Second, all the shapes were oriented vertically and contained a small black dot on the top or bottom, and participants were instructed to respond based on the location of the dot on the target item (see Figure 4). Like in Experiment 1, participants responded with the ‘s’ and ‘l’ keys, but, unlike in Experiment 1, the stimulus-response mapping was counterbalanced across participants. In addition, participants were seated in front of 22-inch LCD display monitors with a resolution of 1680 × 1050 and a refresh rate of 60 Hz, with a viewing distance of approximately 50 cm with the aid of a chin-and-forehead-rest.

**Figure 4:**
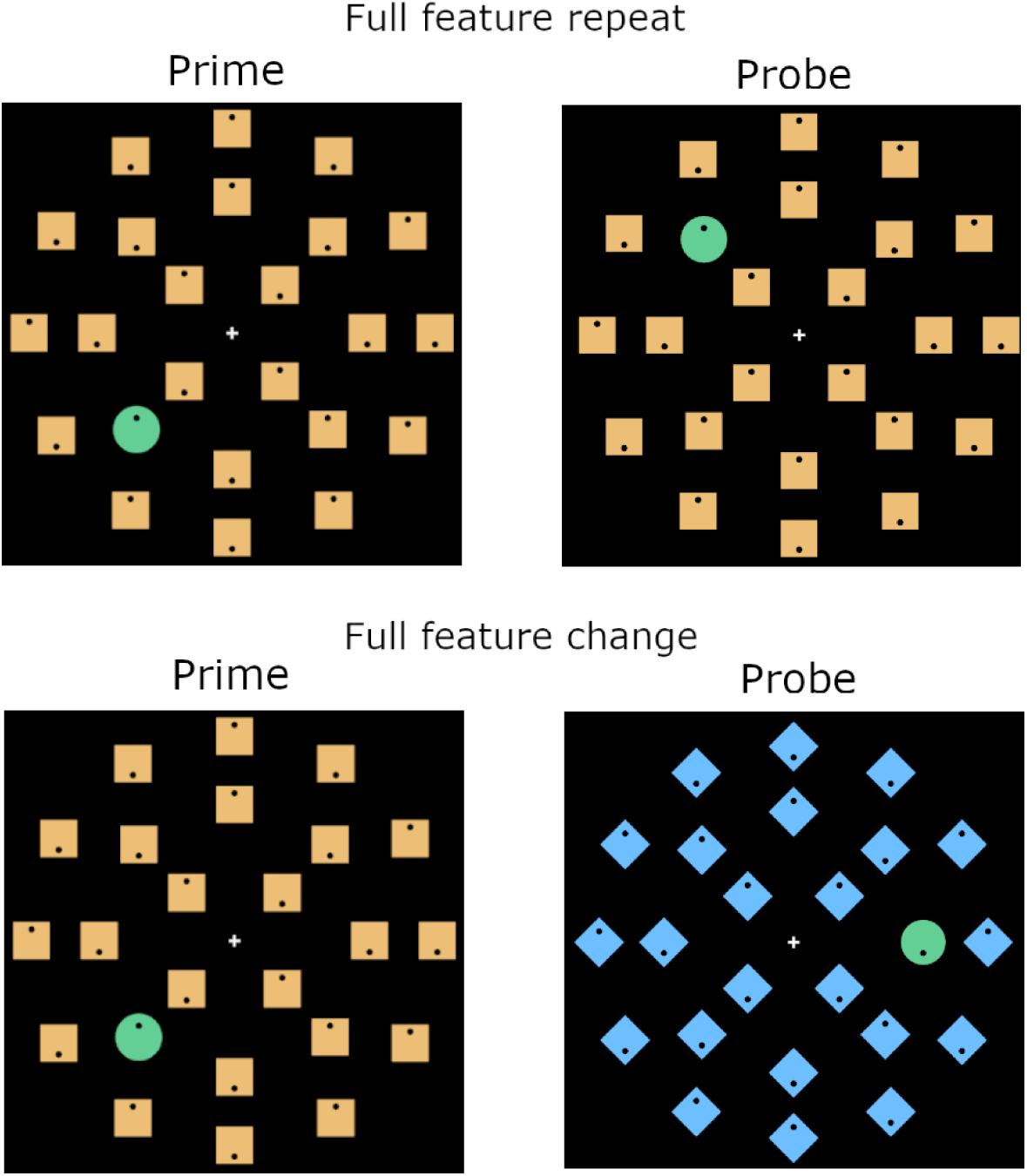
Examples of the search displays used in Experiment 2. In the full feature repeat condition, the non-target color and shape were repeated between the prime and probe trial, while in the full feature change condition they both changed. The task was to locate the target which was always a green circle and respond based on whether the black dot inside the circle appeared at the top or the bottom. The target always differed from the non-targets in terms of both shape and color.

#### Design and procedure

The organization of the experiments into blocks, mini-blocks, and prime-probe pairs, and the number of trials per condition were all the same as in Experiment 1.

### Results

Similar to Experiment 1, we analyzed the response priming effect and the distractor-feature priming using the inverse efficiency score (IES) on probe trials (see the Appendix for separate analyses of response priming on error rates and RTs). We followed the same procedure as Experiment 1, removing probe trials with an error either on the probe trial itself or on the preceding prime trial (7.6 %), and outliers with RTs faster than 200 ms or slower than 3 s (less than 1%).

Figure 5A shows the mean response priming effect as a function of the target position and the distractor-feature repetition. The response priming was positive for all conditions, with an average of 46 ms, which was comparable with the average response priming from Experiment 1 (51 ms in Exp. 1 vs. 46 ms in Exp. 2, *t*(46) = 0.33, *p* = .74). By visual inspection, the response priming was more marked when the distractor features were fully repeated relative to fully changed. A repeated-measures ANOVA with the factors of Distractor Feature (full repeated vs. full changed) and Target Position (fixed vs. random) confirmed that the response priming effect was 16 ms larger for trials with the repeated distractor features, compared to trials with the fully changed distractor features, *F*(1,23) = 6.59, *p* = .017, *η_*p*_*^2^ = .22. Again, this was the test the power-analysis was calculated for.

**Figure 5:**
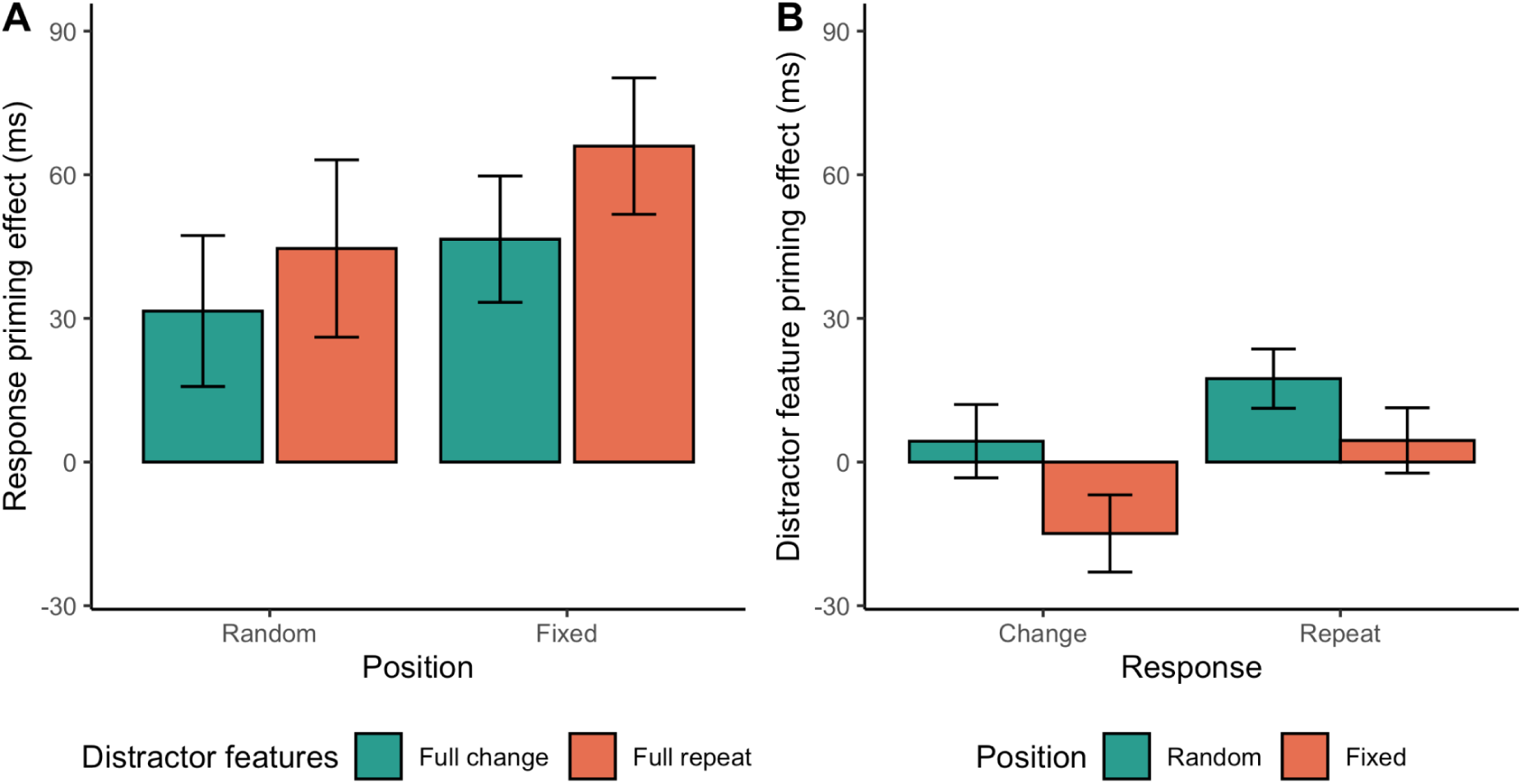
Response priming effect on the inverse efficiency score (IES), i.e. the difference between the IES on response change trials and the IES on response repeat trials, for probe trials from trial pairs where both non-target features where changed (full change) and trial pairs where both features were repeated (full repeat) in Experiment 2. Error bars indicate the standard error of the mean.

However, the factor of the target position was only marginal, *F*(1,23) = 3.48, *p* = .075, *η*_*p*_^2^ = .13. Similar to Experiment 1, further analysis, including the partial repetition (reported in the Appendix), revealed that the response priming was mainly driven by a single distractor feature - color, rather than shape. In the compound task (Figure 4), it appears that the localization of the target was more influenced by color than by shape. Thus, both experiments hint that the response priming is likely contingent on the feature relevance of the task. Taking trials with the partial feature repetition into account, the factor of the Target Position was significant. The response priming was larger when the target position was fixed, compared to trials when the target position was random (*F*(1,23) = 6.42, *p* = .019; see the Appendix). Similar to Experiment 1, the interaction between Target Position and

Distractor-feature Repetition was not significant, *F*(1,23) = 0.22, *p* = .64, *η*_*p*_^2^ = .01. For trials with full-changed distractor features and random target positions, however, the response priming was marginally larger than zero, *t*(23) = 2.04, *p* = .053, indicating that the distractor-response binding (DRB) may not purely account for the response priming we found here.

Figure 5B shows the distractor-feature priming effect. A repeated-measures ANOVA on the distractor feature priming effect revealed a significant effect of response repetition *F*(1,23) = 6.6, *p* = .017, *η*_*p*_^2^ = 0.22, but neither the target position (*F*(1,23) = 3.2, *p* = .09, *η_*p*_*^2^ = 0.12) nor the interaction between response repetition and target position (*F*(1,23) = 0.22, *p* = .64, *η*_*p*_^2^ = 0.09) were significant. The distractor-feature priming was significantly greater than zero (*M* = 11 ms) on trials with repeated responses (*t*(23) = 2.4, *p* = .024, two-tailed), but it did not significantly differ from zero on trials with changed responses (*t*(23) = -1.32, *p* = .20). Just like in Experiment 1, we observed the significant distractor-feature priming only when the response was repeated, providing further evidence that this effect may be a result of retrieval of the correct response when both the distractor features and the response were repeated.

To further identify potential contribution of the response priming when the target position was random, we further examined how the response priming effect depended on how much the target location changed from prime to probe trial^3^. Figure 6 shows the response priming gradually decreased when the prime-probe target distance increased. A repeated-measures ANOVA with the factors of Distractor-feature Repetition and Target Distance on the response priming confirmed a significant effect of Target Distance (*F*(4,88) = 6.34, *p* < .001, *η*_*p*_^2^ = .22). Neither Distractor-feature Repetition, (*F*(1,22) = 3.78, *p* = .07, *η*_*p*_^2^ = .15 nor the interaction between Distractor-feature Repetition and Target Distance (*F*(4,88) = 0.43, *p* = .79, *η*_*p*_^2^

**Figure 6:**
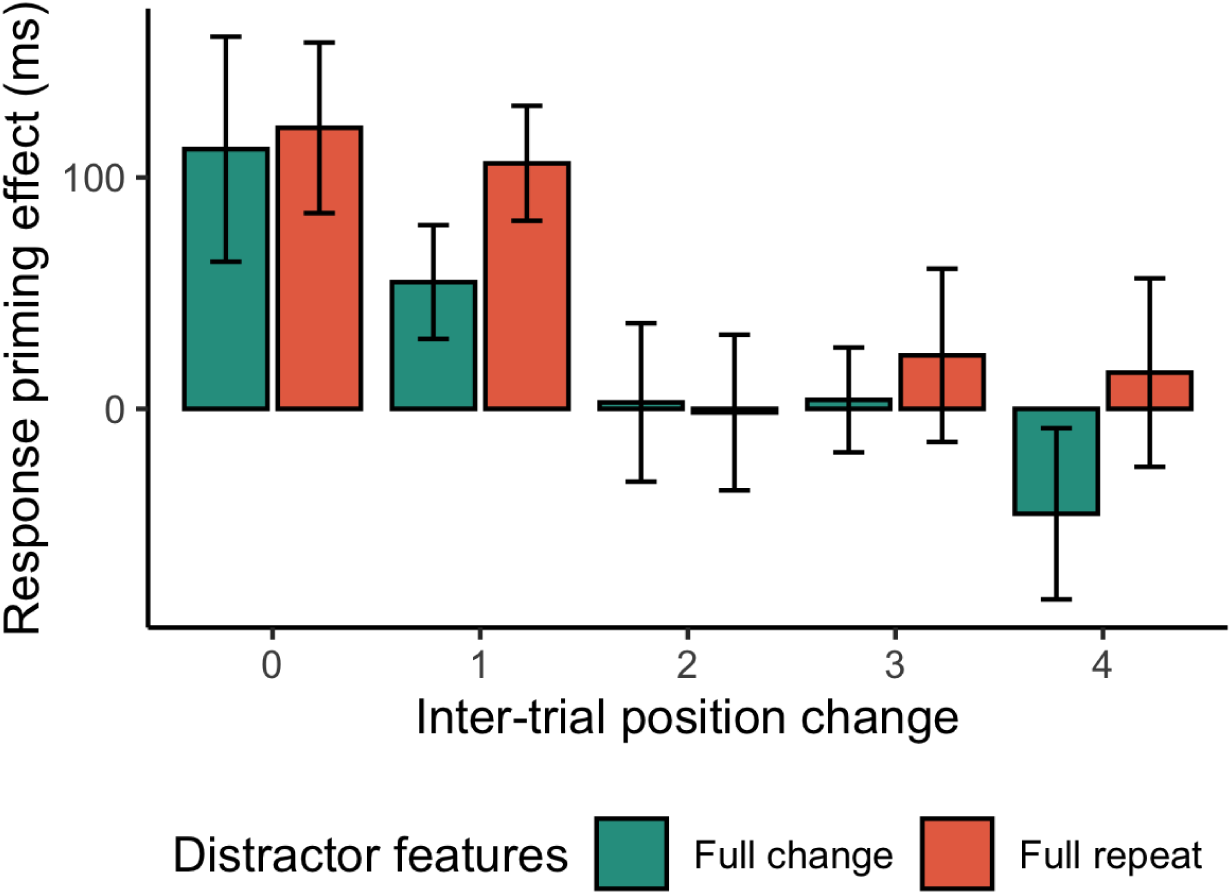
Response priming effect on the inverted efficiency score (IES), in the random target location block of Experiment 2, as a function of the distance the target moved between prime and probe trial (inter-trial position change). Error bars indicate the standard error of the mean.

= .02) was significant. Further post-hoc analyses revealed that the response priming was only significant when the inter-trial position changes was no more than one (dist 0: *t*(22) = 3.43, *p*_bonf_ = .012, dist 1: *t*(22) = 3.89, *p*_bonf_ = .004, all other p > .6).

### Discussion

Instead of using basic feature orientation, Experiment 2 used the compound feature - the location of the dot - as the response-defining feature to reduce the potential adoption of the target-feature search strategy. Here we replicated the findings of Experiment 1. The response priming was comparable between Experiments 1 and 2. Importantly, we showed that both the distractor-feature repetition and the target position repetition contributed to the response priming. Again, we showed the distractor-feature priming was relatively weak, mainly contributed by those trials when the response was repeated. Different from Experiment 1, we found that the response priming effect was manifested when the target position was repeated or the neighbour of the previous target position for both the distractor features fully changed and fully repeated conditions, suggesting target-position-response binding is also a key factor contributing to the response priming.

## General discussion

The present study set out to investigate distractor-response binding (DRB) in visual search, a topic that has garnered limited focus within the search community. We designed two experiments, maintaining constant target features throughout to minimize reliance on any target-related dimension-weighting search strategies. The distractor features, however, varied across the prime-probe pairs. Our findings revealed that response priming occurred in both experiments, with a greater effect when the distractor features were repeated, compared to when they were changed, and when the target position was fixed, as opposed to when it varied. Interestingly, our comprehensive analyses, including trials with both full and partial feature repetition (detailed in the Appendix), revealed that response priming was primarily driven by a specific distractor feature (shape in Experiment 1 and color in Experiment 2). This suggests that participants might have concentrated on the most distinctive and task-relevant feature (color or shape) differentiating the target from distractors and down-weighted the other feature. Additionally, when the target’s position was fixed, response priming was also enhanced. These findings suggest that response binding plays a critical role in the absence of the target-related up-weighting strategy.

Distractor-response binding (DRB) is a concept well-documented in the action control literature (e.g. Laub and Frings 2020; Singh and Frings 2020). In a typical DRB task, an array of letters appeared and participants are instructed to focus on the target, identified by its color and position, while ignoring the flanking distractor letters (e.g., Frings, Rothermund, and Wentura 2007). The response binding effect has been interpreted in the framework of binding and retrieval in action control (BRAC, Frings et al. 2020). According to BRAC, features of the stimulus environment, the response in that environment, and its subsequent effects are collectively integrated into an event-file (Hommel 2004). Whenever attributes of the event file recur, it triggers the retrieval of that specific event file, which in turn affects performance. The repetition of the distractor features, even if they are task-irrelevant, can reactivate the previous associated event-file, including the response. This reactivation enhances performance if the retrieved response is compatible with the currently demanded response. Until now, the DRB effect has mainly been examined in non-search flanker tasks. Here, our study extends this understanding, showing the DRB effect was also present in visual search tasks. Yet, our findings of the distractor-response binding (DRB) effect in visual search do not just extend the DRB previously observed mainly in the flanker task. The main difference between the two tasks is that the visual search involves a search stage. Our study found that the DRB effects only occurred when the target position was repeated or near the previous target position, which was not apparent in the flanker task due to the absence of uncertainty in the target position. Our findings highlight the important role of the target position, a critical factor that has been largely overlooked in the previous action control literature.

Previous studies have identified response priming as the “secondary” feature in visual search (Zehetleitner, Rangelov, and Müller 2012; V. Maljkovic and Nakayama 1994; Yashar and Lamy 2011; Töllner et al. 2008). For instance, Lamy et al. (2011) distinguished between the target feature repetition and response repetition by mapping four second features to two alternative responses (i.e., two of which were mapped to one response). They showed that the response-based component of feature priming effect could be used, but is not mandatory, when the task difficulty is high. Zehetleinter and colleagues (2012) also observed response priming resulting from the interaction between changes in a task-irrelevant feature in the target and the preparation of a response. Some researchers believe that response-based feature priming is a result of an episodic memory retrieval mechanism (Lamy, Zivony, and Yashar 2011; Huang, Holcombe, and Pashler 2004; Ásgeirsson and Kristjánsson 2011), which is in general compatible with the BRAC framework. However, previous studies on response priming in visual search have mainly focused on the target-related or response-related features, neglecting distractor-response binding. In the present study, the target features were kept constant throughout the experiment, meaning the response-based secondary feature was fixed. Despite this, we found that the distractor-feature repetition also contributed to the response priming, suggesting that the response binding is broader than the target-related feature priming.

Both experiments revealed that repeating the distractor feature facilitated search performance, confirming the distractor feature priming and replicating previous research (Lamy et al. 2008). Interestingly, though, our study showed that this priming was weaker than the response priming (only about half the magnitude) and only occurred when the response was repeated. Essentially, the distractor feature priming we observed can be interpreted by the response priming, as the feature priming in the baseline of response-change was not significant. In this aspect, the BRAC framework may have broader prediction for intertrial priming. It is important to note that the BRAC framework complements theories developed in visual search, such as the dimension-weighting account (DWA, Müller, Heller, and Ziegler 1995). The DWA emphasizes the importance of up-weighting target-relevant dimensions while the BRAC highlights the relevance of the retrieval of encoded event-files. In the present study, we limited changes of the target-related features that related to DWA to highlight the effects of distractor-response binding (DRB). Nevertheless, the BRAC framework can be enhanced by weighting aspects as described by the DWA. In fact, while several previous studies refer to weighting in the context of event-coding, the precise mechanism is not understood (or even discussed) so far. Connecting visual search (that has a rich background on feature weighting processes) with BRAC moves the action control research forward. Feature weighting might be modeled as described by DWA even in non-search contexts.

In addition, in both experiments, we showed a robust target position priming. The response priming was larger when the target was repeated at the same position, similar to previous research on target position priming (Allenmark et al. 2021). The response priming was mainly contributed by those trials with the prime-probe target distance not exceeding one location. That is, the response priming effect was only manifestable when the target remained roughly in the same or neighboring position (within 4.6° of visual angle). According to the BRAC framework (Frings et al. 2020), response priming is a result from retrieval and activation of previous event-file. The fact that the response priming was contingent on the target position suggests that the response-binding retrieval likely occurs during the late stage of target identification or response selection or both. Otherwise, the predominant distractor features repetition (almost the whole display) might well activate the previous even-file during the preattentive and early search stage, resulting in a general response priming effect, independent of the target position. Our results, however, reject this possibility. Our findings are consistent with previous studies that have shown that the response repetition effect operates at a late post-perceptual response-related stage (Yashar and Lamy 2011; Lamy, Yashar, and Ruderman 2010; Zehetleitner, Rangelov, and Müller 2012).

Our paper can be seen as a first step towards a more holistic understanding of human action. In fact, we pinpoint very specific processes relevant to human action in experimental paradigms by controlling as many influences as possible that are not directly relevant to the specific task at hand (e.g. no search in a typical flanker task). While this approach led to many intriguing insights into human cognition and action it also has limitations. In particular, sometimes the ‘big picture’ gets lost - that is, we do not want to explain visual search or binding processes in a particular task but ultimately human behavior in general. For our paper that means that we do want to acknowledge the influences of binding and retrieval in visual search paradigms on the one hand and of visual search and location aspects for action on the other hand. Still, we used typical experimental paradigms here (that are far from real world behavior). Yet, by linking these we at least approach understanding human behavior from different perspectives and combine search and action aspects.

In summary, the present results extend the scope of the BRAC framework to response priming in intertrial priming in visual search and show that distractor feature repetition modulates intertrial priming through binding and retrieval. It is also noteworthy that we have shown that distractor-response binding can extend to visual search, and that response binding depends on the position of the target stimulus, which likely occurs in the late phase of target identification and response selection. Binding and retrieval mechanisms are at work, even if researchers did not intend to analyze these processes (Henson et al., 2014). However, especially when researchers use sequential paradigms, observed effects should probably always be interpreted in terms of these processes-regardless of whether the original research question originated in the action control literature.

## Appendix: Comprehensive analyses of reaction times, error rates, and inverse efficiency scores

This appendix contains comprehensive analyses of reaction times (RTs), error rates, and inverse efficiency scores (IESs), including the effects of partial and full repetitions. The findings detailed below reinforce the conclusions drawn in the main manuscript, where it was established that response priming effects were more pronounced when distractor features were full repeated.

### Experiment 1

Figures A1 (A and C) show the average RTs and error rates as a function of for the distractor shape and color changes, separated for the target position (change vs. repeat) in Experiment 1, while Figure A1 (B and D) depict the corresponding response priming effects (calculated as the difference between the average RT or error rate on response change vs. repeat trials).

**Figure A1:**
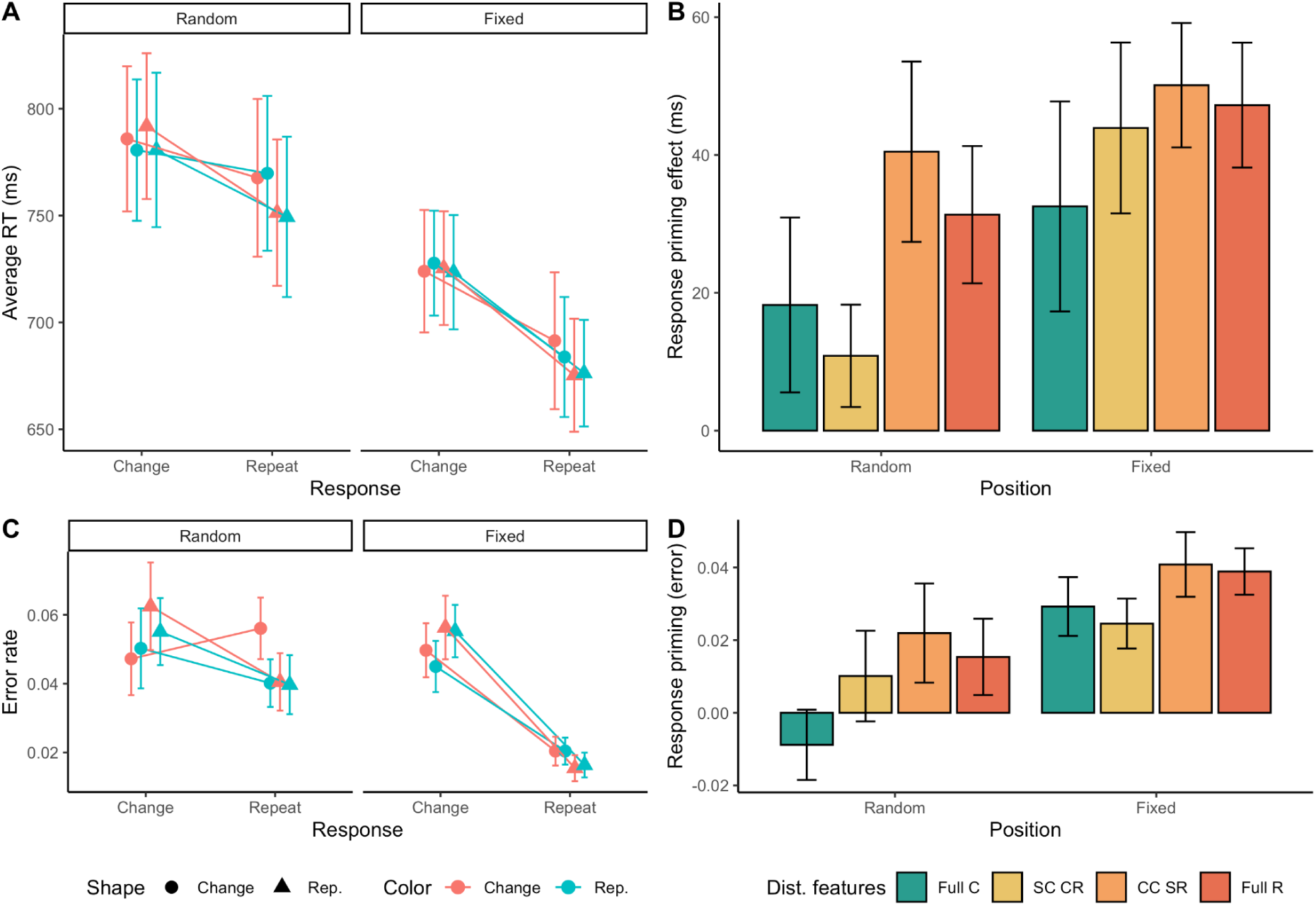
Mean reaction times (A) and error rates (C) and response priming effects on RTs (B) and error rates (D) in Experiment 1in the different color and shape repetition conditions: full change (Full C), shape change and color repeat (SC CR), color change and shape repeat (CC SR) and full repeat (Full R). Error bars indicate the standard errors of the correspondent mean.

A repeated-measures ANOVA on RTs with Response Repetition (repeat vs change), Distractor Color Repetition (repeat vs change), Distractor Shape Repetition (repeat vs change) and Target Position (fixed target location vs. random target location) as factors revealed significant main effects of Response Repetition (*F*(1,23) = 35.98, *p* < .001), Distractor Shape Repetition (*F*(1,23) = 5.69, *p* = .026) and Target Position (*F*(1,23) = 32.30, *p* < .001). RTs were significantly faster when either the response or the shape was repeated and in the fixed target location compared to the random target location block. In addition, there were significant interactions between response repetition and shape repetition (*F*(1,23) = 4.82, *p* = .039) and between response repetition and target position (*F*(1,23) = 4.84, *p* = .038). The response priming effect was significantly larger when distractor shape was repeated compared to when it changed and in the fixed target location block compared to the random target location block.

A similar pattern emerged for the error rates, with significant main effects of Response Repetition (*F*(1,23) = 11.39, *p* = .003) and Target Position (*F*(1,23) = 8.83, *p* = .007), along with significant interactions between Response Repetition and Shape Repetition (*F*(1,23) = 13.87, *p* = .001), and between Response Repetition and Target Position (*F*(1,23) = 12.84, *p* = .002). Thus, the response priming effects on both RTs and error rates were influenced similarly by context repetition; both effects were smaller when the target position was not repeated and when the distractor shape was not repeated.

For completeness we also analyzed the response priming effect on the inverse efficiency score across all conditions (see figure A2). Consistent with the analysis of reaction times above, there were significant main effects of Distractor Shape Reptition (*F*(1,23) = 16.5, *p* < .001) and Target Position (*F*(1,23) = 9.0, *p* = .006), with no other effects showing significance (all *p*s > .3).

**Figure A2.**
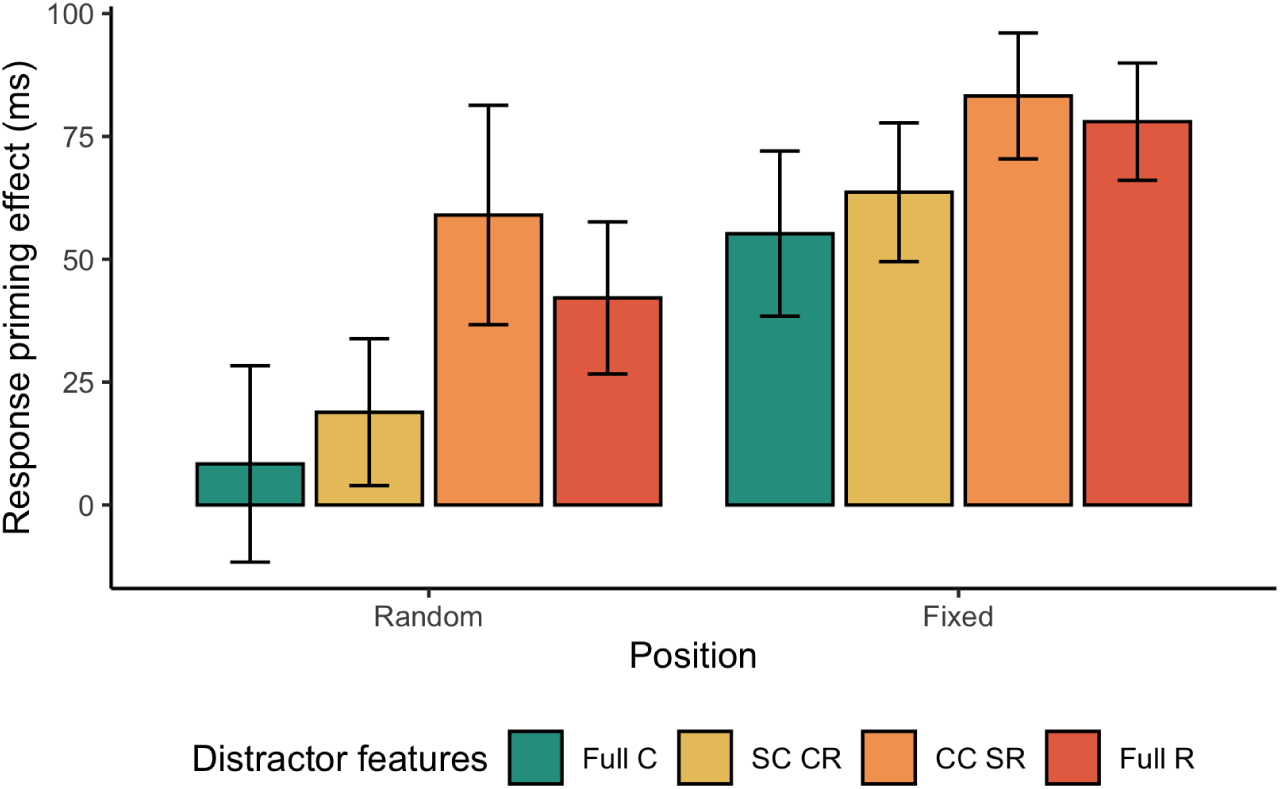
Response priming effects calculated from inverse efficiency scores (IES) in Experiment 1, across various color and shape repetition: full change (Full C), shape change and color repeat (SC CR), color change and shape repeat (CC SR) and full repeat (Full R). Error bars indicate the standard error of the mean.

These results are consistent with the results based on the IESs in the main manuscript and further indicate that in Experiment 1 the distractor shape had a larger influence on response priming than the distractor color.

### Experiment 2

Figure A3 shows the average RTs and error rates across various conditions in Experiment 2, along with the corresponding response priming effects (calculated as the difference between the average RTs or error rates in response change trials and the those in response repetition trials).

**Figure A3:**
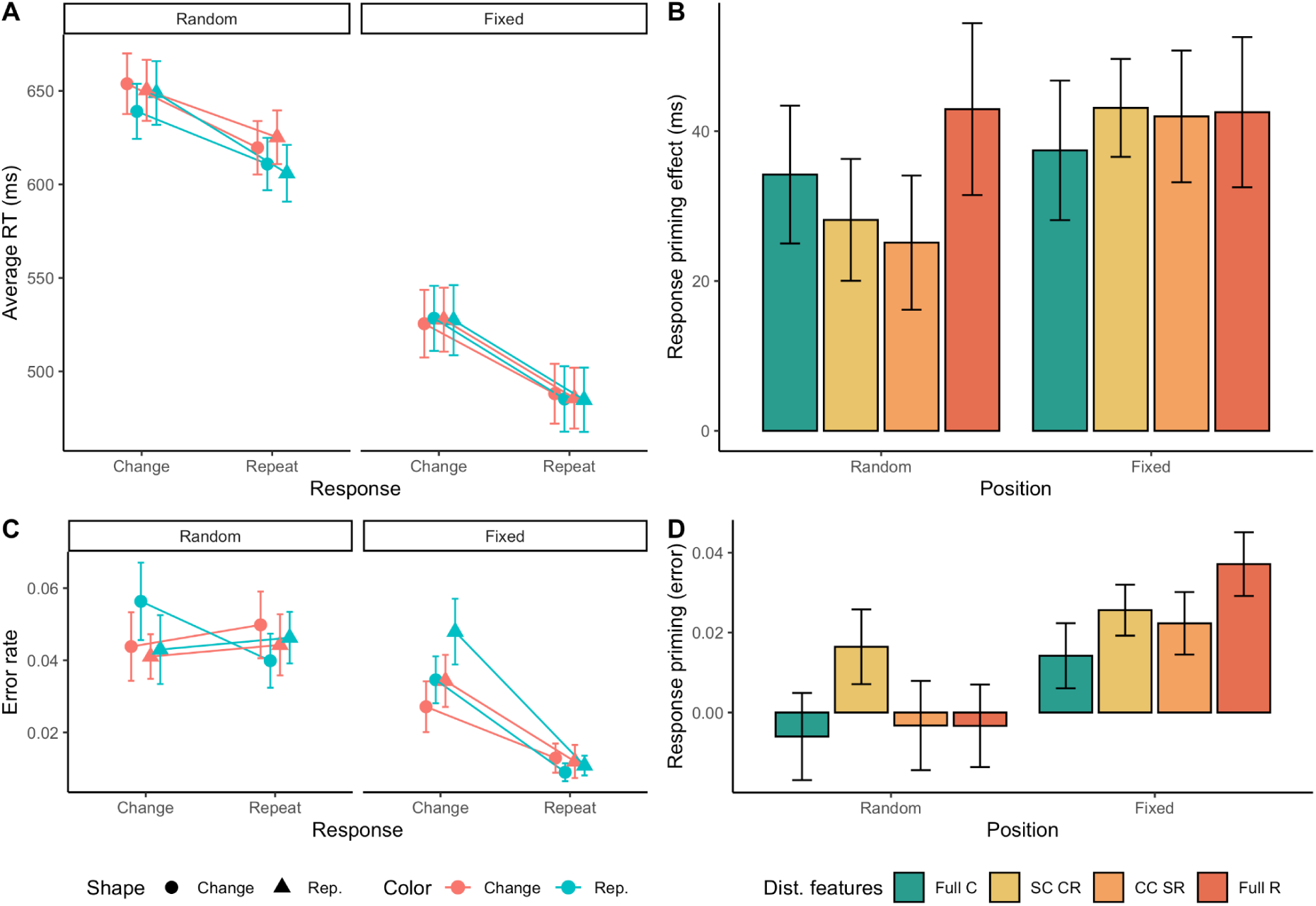
Mean reaction times (A) and error rates (C) and response priming effects on RTs (B) and error rates (D) in Experiment 2under different color and shape repetition conditions: full change (Full C), shape change and color repeat (SC CR), color change and shape repeat (CC SR) and full repeat (Full R). Error bars indicate the standard error of the mean.

A repeated measures ANOVA on RTs with Response Repetition (repeat vs change), Distractor Color Repetition (repeat vs change), Distractor Shape Repetition (repeat vs change) and Target Position (fixed target location vs. random target location) as factors revealed significant main effects of Response Repetition (*F*(1,23) = 23.81, *p* < .001), Distractor Color Repetition (*F*(1,23) = 8.31, *p* < .01) and Target Position (*F*(1,23) = 19.03, *p* < .001). RTs were significantly faster when either the response or the distractor color was repeated and in the fixed target location compared to the random target location block. In addition, there were significant interactions between Response Repetition and Target Position (*F*(1,23) = 6.42, *p* = .019) and between Distractor Color Repetition and Target Position (*F*(1,23) = 9.19, *p* = .006); the response priming effect was significantly larger in the fixed target location block compared to the random target location block while the color repetition effect was larger in the random location block. For the error rates there was a significant main effects of Response Repetition (*F*(1,23) = 5.01, *p* = .035) and Target Position (*F*(1,23) = 47.06, *p* < .001) and significant interactions between Response Repetion and Distractor Color Repetition (*F*(1,23) = 7.12, *p* = .014) and between Response Repetition and Target Position (*F*(1,23) = 14.43, *p* < .001) as well as a significant three-way interaction between Response Repetition, Distractor Shape Repetition and Target Position (*F*(1,23) = 5.21, *p* = .032).

In summary, Experiment 2, like Experiment 1, showed that response priming effects in terms of both RTs and error rates were substantial larger in the blocks where the target position was fixed. However, in contrast to Experiment 1, only the response priming effect on error rates was significantly influenced by the context, and it was more affected by the distractor color than by the distractor shape.

We again analyzed also the response priming effect on the inverse efficiency score across all conditions (see figure A4).

**Figure A4.**
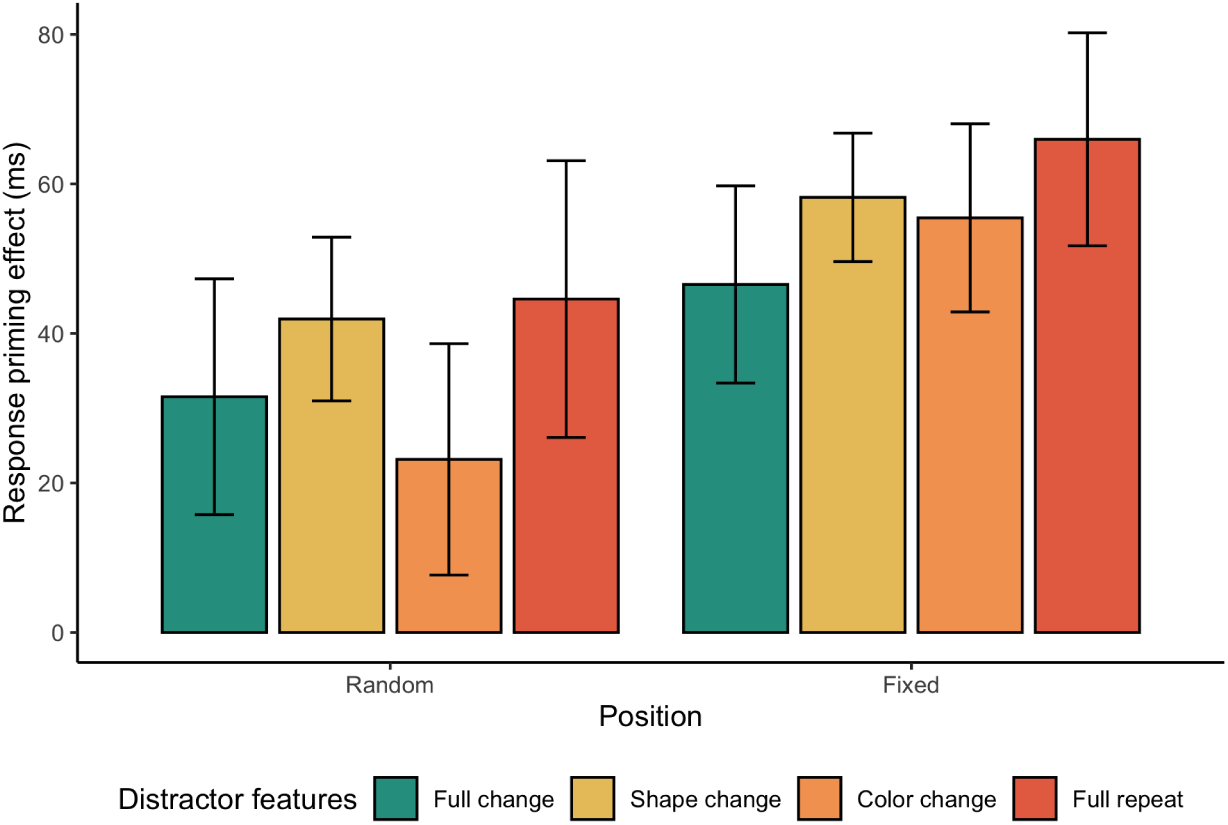
Response priming effects on inverse efficiency scores in Experiment 2 in the different color and shape repetition conditions: full change, shape change and color repeat (SC CR), color change and shape repeat (CC SR) and full repeat. Error bars indicate the standard error of the mean.

Consistent with the analysis of reaction times above, there were significant main effects of Distractor Color Repetition (*F*(1,23) = 8.7, *p* = .007) and Target Position (*F*(1,23) = 9.0, *p* = .006). No other effects were significant (all *p*s > .1).

In summary, the full analyses reported in this appendix revealed nuanced aspects of our findings. In both experiments, response priming decreased when distractor features changed compared to when they were repeated. However, the key influencing factor varied between the experiments: in Experiment 1, it was primarily the repetition versus change of the distractor shape that impacted response priming, while in Experiment 2, the critical factor was the repetition versus change of the distractor color. It should be noted that the task of Experiment 1 was orientation discrimination, while the task in Experiment 2 was to discriminate the location of a small dot inside the target. The contribution to the response priming may hinge on the task relevance and saliency of the feature.

## Footnotes

1 In principle the DWA framework would allow for an element of feature-specificity in attentional selection over and above dimension-specificity, as observed by Found and Müller (1996) and Müller et al. (2003) especially for color-defined targets. For instance, entry-level coding of a particular target feature might be enhanced top-down by setting up the appropriate template, giving this feature an edge (see a similar note by Tsai et al. 2023).

2 While the Multi-weighting system (MWS) would also predict the repetition benefit of the response-defining features (Zehetleitner, Rangelov, and Müller 2012; Rangelov, Müller, and Zehetleitner 2012), DRB incorporates not only the target features but also the distractor features. This makes DRB a more versatile account for analyzing and understanding inter-trial priming.

3 For this analysis one participant was excluded due to not having any valid trials in one of the conditions after removing error trials and outliers.

